# *In-vitro* Reconstitution of Membrane Contact Sites between the Endoplasmic Reticulum and Lipid Droplets

**DOI:** 10.1101/2021.06.07.447315

**Authors:** Sukrut Kamerkar, Jagjeet Singh, Subham Tripathy, Hemangi Bhonsle, Mukesh Kumar, Roop Mallik

**Affiliations:** Department of Biosciences and Bioengineering, Indian Institute of Technology Bombay, Mumbai 400076, India; Department of Biological Sciences, Tata Institute of Fundamental Research, Mumbai 400005, India

**Author notes:** Co-first authors (Sukrut Kamerkar, Jagjeet Singh).

## Abstract

Coordinated cell function requires inter-organelle communication across Membrane Contact Sites (MCS). Here we deposit ER-enriched microsomes purified from rat liver or from cultured cells on a coverslip in the form of a continuous planar membrane. We visualize real-time protein and lipid exchanges across MCS that form between this ER-mimicking membrane and lipid droplets purified from rat liver. An Optical trap is used to demonstrate physical tethering of individual lipid droplets to the ER-mimicking membrane at MCS, and to directly measure the strength of this tether. *In-vitro* MCS formation changes dramatically in response to metabolic state and immune activation in the animal. Surprisingly, we find that the Rab18 GTPase and Phosphatidic acid are common molecular factors to control both of these pathways. This assay could possibly be adapted to interrogate MCS formation between other membranes (e.g. mitochondria, peroxisomes, endosomes etc.), and abnormalities therein that cause neurological, metabolic and pathogenic diseases.

---

Lipid droplets (LDs) are triglyceride-rich cellular organelles that act as an energy-depot and lipid reservoir for membrane biosynthesis(1–3). LDs are also dynamic, exchanging proteins and lipids with the endoplasmic reticulum (ER), mitochondria and other organelles via formation of MCS(3–5). LD-ER contacts are a topic of intense discussion because of their role in neurological and metabolic disorders, pathogen infection and hepatic steatosis(3). Unfortunately, the molecular machinery acting at these sub-microscopic contacts inside cells remain poorly understood and difficult to interrogate. Synthetic droplet emulsified vesicles replicate certain biophysical aspects of LD-ER contacts(6), but they are devoid of the attendant cellular machinery. We therefore wondered if ER-derived microsomal vesicles could be deposited onto a coverslip as a continuous planar membrane that mimics the ER, and if purified cellular organelles (e.g. LDs) could form MCS with this ER-mimicking membrane (hereafter called the **<u>m</u>**icrosomal **<u>S</u>**upported **<u>L</u>**ipid **<u>B</u>**ilayer; **mSLB**). Such an assay could allow controlled *in-vitro* manipulation and visualization of inter-organelle contacts, also shedding light on the molecular mechanisms behind two recent discoveries:-**(i)** Dramatic changes in LD-ER interactions inside the liver across fed/fasted states(7, 8) to facilitate systemic lipid homeostasis in a manner relevant to fatty liver disease(9), and **(ii)** A function for LDs in the liver as innate immune hubs to protect against bacterial infection(10).

Accordingly, we modified the conventional technique(11) of preparing planar supported lipid bilayers (SLBs) by replacing synthetic liposomes with microsomal vesicles that were prepared from rat liver or from cultured cells. Established ER markers such as Calnexin (Fig S1:A) and S6 (Fig S1:B) were detected on the mSLB by antibody staining. Purity of microsomal preparations was confirmed as described earlier(7, 8) (Fig S1:C). We chose BODIPY-C12 as a direct probe to visualize the mSLB because this fluorescent fatty acid analog incorporates spontaneously into membranes. Microsomes were first doped with trace amounts of BODIPY-C12 and then introduced into a flow cell containing a hydrophilic plasma-cleaned coverslip. Excess microsomes were washed off, and the mSLB was imaged (Fig 1A). Fluorescence recovery after photobleaching (FRAP) of BODIPY-C12 revealed the mSLB as a continuous membrane with high mobility of lipids within the mSLB (Fig S1:D,E), as also reported for artificial membrane tubes(12). The diffusion coefficient of BODIPY-C12 in mSLBs (=2.3±0.17μm2sec^-1^) was lower than in protein-free conventional SLBs prepared using phosphatidylcholine containing liposomes (=3.0±0.32μm2sec^-1^; Fig S1:F,G,H), likely because ER-associated proteins act as a diffusion barrier.

**Figure 1.**
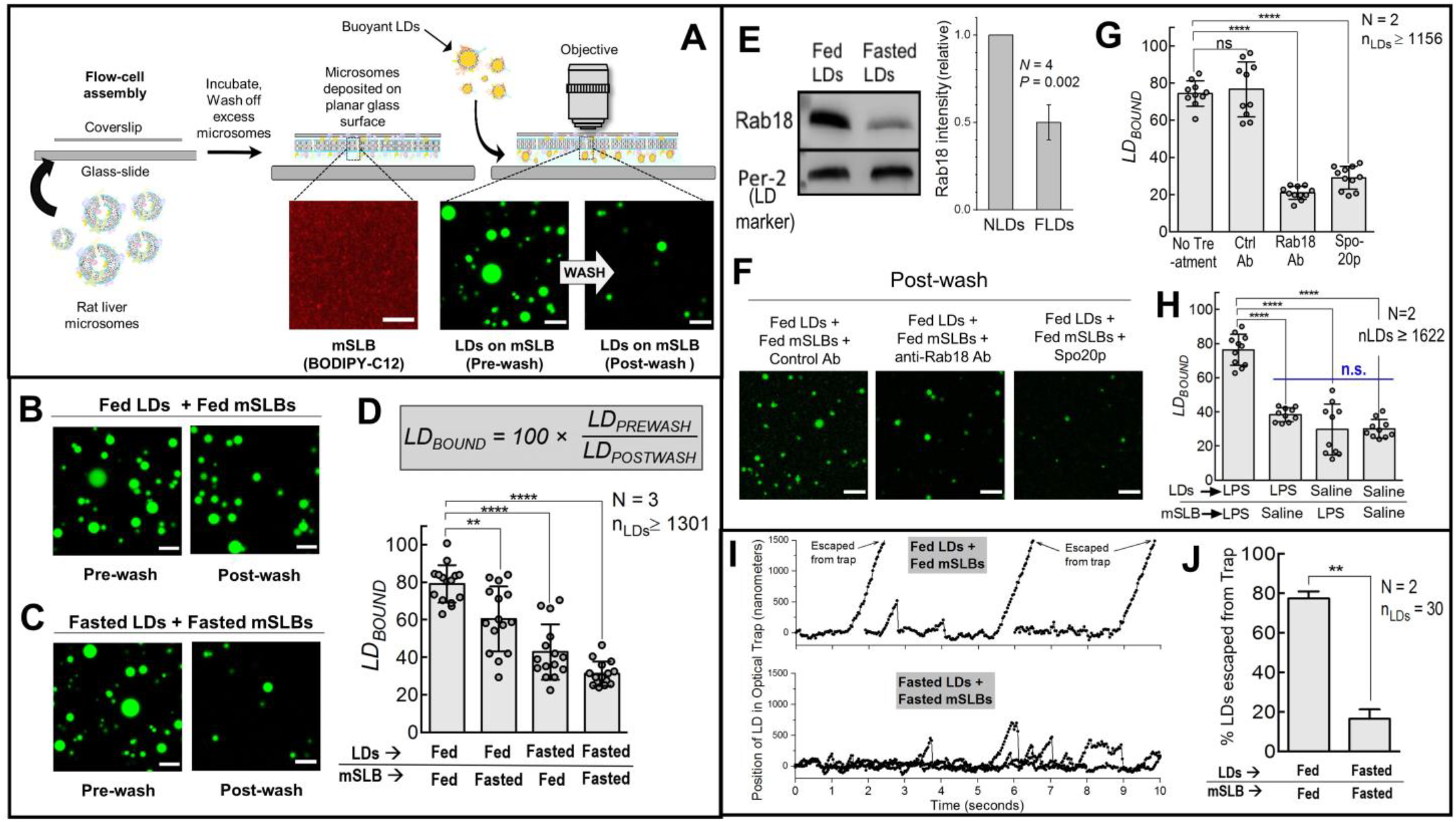
*In vitro* Reconstitution of LD-ER membrane contact sites. A. Schematic depicting formation of ER-mimicking <u>m</u>icrosomal <u>S</u>upported <u>L</u>ipid <u>B</u>ilayer (mSLB). Rat liver microsomes are prepared from rat liver, sonicated and introduced inside a chamber containing a plasma-cleaned coverslip. Spontaneous fusion of microsomes to the coverslip leads to formation of a continuous mSLB. Excess microsomes are washed off and LDs are introduced into the chamber. Due to their buoyancy, LDs float and come in contact with mSLBs and a fraction of them bind to the mSLB (presumably via MCS). Unbound LDs are washed off after incubation (see pre-wash and postwash images). Scale bar = 10 μm. Bodipy-C12 labelled mSLB shown in red pseudo-colour. B. Representative images of fed-LDs (green) incubated with fed-mSLB. Pre-wash and post-wash images are shown. Scale bar = 10 μm. C. Representative images of fasted-LDs (green) incubated with fasted-mSLB. Pre-wash and post-wash images are shown. Scale bar = 10 μm. D. Quantitation of *LD_BOUND_* under different metabolic states. Each point on the graph represents a field of view with ~100 LDs. Data represent mean ± SD. N represents number of independent pair of animals used in the experiments, n_LDs_ represents the total number of LDs observed. **** *p* < 0.0001, ** *p* = 0.003 using Mann–Whitney test. E. Immunoblot for Rab18 and perilipin-2 (loading control) on fed and fasted LDs isolated from rat liver. Panel on right quantifies Rab18 levels on LDs. Rab18 intensity on fed-LDs was always taken = 1. Result of one-sample *t*-test is shown across four independent experiments. F. Representative images of post-wash fed-LDs added to fed-mSLB treated with anti-FLAG-IgG Ab (control), anti-Rab18 antibody or Spo20p protein. LDs are labeled in green. Scale bar = 10 μm. G. *LD_BOUND_* calculated for fed-LDs bound to fed-mSLB under conditions mentioned in (F). Each point on the graph represents a field of view with ~100 LDs. Data represent mean ± S.D. *N* represents the number of independent pair of animals used. n_LDs_ represents the total number of LDs observed. **** *p* < 0.0001; ns = not significant, Mann–Whitney test. H. *LD_BOUND_* for LDs isolated from liver of fasted rats that had been injected with saline (control) or LPS. These LDs were incubated with mSLB prepared from liver of fasted rats that had been injected with saline (control) or LPS in four possible combinations. Data represent mean ± S.D. *N* represents the number of independent pair of animals used, n_LDs_ represents the total number of LDs observed. ** *p* < 0.01, Mann–Whitney test. I. Position-time plot for fed-LDs and fasted-LDs held in an optical trap on respective mSLB surfaces. Displacement of LDs as a function of time was obtained by video tracking while moving the mSLB with a piezo stage. J. Percent LDs escaped from the optical trap. ** *p* < 0.01, Mann–Whitney test.

We next prepared LDs from the liver of normally fed rats (fed-LDs) and verified their purity as described(7, 8, 13) (Fig S1:C). Fed-LDs were added to mSLBs prepared from liver of fed rat (fed-mSLBs) followed by incubation in the orientation shown in Fig 1A (coverslip on top; see schematic in Fig 1A). This geometry allows the buoyant LDs to float up and come in contact with the mSLB on the coverslip. We then imaged the flow-cell looking down through an **<u>*upright*</u>** confocal microscope, LDs were imaged right after the incubation, the flow-cell was washed to remove the unbound LDs, and remaining LDs were imaged again (Fig 1A). LDs still retained on the mSLB post-wash are assumed to have stable LD-ER contacts. LD numbers were counted in pre-wash (=*LD_PREWASH_*) and post-wash (=*LD_POSTWASH_*) images. We then calculated *LD_BOUND_* (=100 × *LD_POSTWASH_/LD_PREWASH_*) to measure how efficiently potential MCS form between LDs and the mSLB. Upon flowing in a dilution series of LDs onto mSLBs we expectedly found that *LD_PREWASH_* reduces with dilution (Fig S2:A), but *LD_BOUND_* was unchanged (Fig S2:B). The use of this ratio (*LD_BOUND_*) therefore makes our estimate of LD-ER contact formation largely independent of variation in LD numbers across samples. We therefore expect that the quantity *LD_BOUND_* reflects changes in LD-ER binding affinity arising from changes in LD/ER membrane composition, for example in response to metabolic transitions(7–9) or immune activation(10).

We next used rats fasted for 16 hours to prepare LDs (fasted-LDs) and mSLBs (fasted-mSLBs) from the liver(7, 8). Fed-LD and fasted-LD samples were adjusted to equal optical density (via dilution) to ensure that they had approximately equal number of LDs per unit volume(7, 8). This “normalization” was further confirmed by measuring triglyceride content (Fig S2:D). As expected because of the earlier normalization step, *LD_PREWASH_* was comparable between fed-LD and fasted-LD samples (Fig S2:C). Note that fed-LDs and fasted-LDs have similar size distribution (Fig S2:E), as reported earlier(13). Next, fed-LDs or fasted-LDs were introduced over fed-mSLBs or fasted-mSLBs in four possible combinations and imaged (Fig 1B, 1C; only two combinations shown). Significantly more fed-LDs remained bound to fed-mSLBs after the wash (*LD_BOUND_* ~80%; Fig 1D), supporting that LD-ER contacts are enhanced in fed state to supply triglyceride for VLDL assembly in hepatocytes(7, 8). The graded reduction of *LD_BOUND_* when moving from fed-fed to fasted-fasted combination (Fig 1D) suggests that binding partners act on both (LD and ER) membranes to sustain MCS in fed state. We next investigated the identity of such molecules.

The Rab18 GTPase associates with NRZ-SNARE complex to form LD-ER contacts(14, 15). This association and LD-ER contacts are important for Dengue and Hepatitis-C virus replication(3). Mutations in Rab18 also cause neurodevelopmental disorders such as the Warburg Micro syndrome(3). Because loss of Rab18 also impairs LD catabolism(16), Rab18 could possibly aid LD-ER contact formation in fed state to catabolize LDs for VLDL production. Indeed, Rab18 levels were dramatically enhanced on fed-LDs compared to fasted-LDs (Fig 1E). Fed-LD to fed-mSLB contacts were inhibited specifically by a Rab18 antibody (Figs 1F, 1G) suggesting that Rab18 may facilitate ER-LD MCS formation in hepatocytes in fed state. To investigate further, mSLBs were prepared from microsomes of cells overexpressing constitutively active (Q67L; GTP-mimic) or inactive (S22N;GDP-mimic) Rab18(15). Both forms of overexpressed protein were equally abundant on microsomes (Figs S2:F and S2:G). When fed-LDs were added to these mSLBs, *LD_BOUND_* was significantly higher for the Rab18 GTP mimic (Fig S2:H). Therefore, LD-ER contacts are likely promoted by GTP-bound Rab18 on the ER.

Phosphatidic acid (PA) recruits the kinesin motor to LDs by directly binding kinesin’s tail domain(7, 8). The yeast SNARE protein Spo20p binds to PA with high affinity, and therefore competitively displaces other PA-bound proteins (e.g. kinesin) from the LD membrane(8). Spo20p-GST reduced LD-mSLB contacts significantly (Fig 1F and 1G), suggesting that MCS formation requires factors that bind to PA on LDs. LDs were recently found to make physical contacts with bacteria to deliver anti-bacterial proteins in response to immune activation(10). Notably, immune activation also appears to promote intimate LD-ER contacts in the liver (Supplementary Figure 6 in Ref.(10)), although this aspect was not explored in that manuscript(10). The exciting possibility therefore emerges that LDs may actually acquire anti-bacterial proteins in response to immune activation from the ER (via ER-LD MCS formation). We therefore activated innate immunity in fasted rats by lipopolysaccharide (LPS) injection(10), followed by LD and mSLB preparation. LDs and mSLB were also prepared from fasted rats injected with saline as a control. We then assayed these LDs and mSLB in four possible combinations. *LD_BOUND_* was highest when both LDs and mSLB were prepared from LPS-treated animals (Fig 1H and Fig S3:A), suggesting the requirement of binding partners on both (LD and ER) membranes for MCS formation. Which molecules may engineer LD-ER contacts in response to immune activation? Immune activation recruits Rab18 to LDs(10), reminiscent of increased Rab18 on fed-LDs (Fig 1E). Further, PA was also enriched on LPS-treated LDs (Fig S3:B-C) reminiscent to increased PA on fed-LDs(8). It is therefore possible that a common molecular pathway involving Rab18 and PA mediates ER-LD MCS formation in response to feeding as well as immune activation.

To verify directly the physical tethering of LDs to the mSLB, individual LDs were held in an optical trap(13). We then moved the mSLB underneath a trapped LD, expecting that binding to mSLB would displace the LD from the trap center (Schematic in Fig S3:D). Fig 1I shows such displacements for fed-LDs on fed-mSLBs, and for fasted-LDs on fasted-mSLBs. In the fed case, most LDs attached to the moving mSLB and escaped from the trap (displacement > 1000nm) within 20 seconds (Supp Movie 1). However, in the fasted case most LDs exhibited transient attachments/detachments without escapes, revealing weaker LD-mSLB interactions (Fig 1I; Supp Movie 2). The optical trap works as a spring of certain stiffness. We have developed a method for calibrating this stiffness for LDs(13), wherein we had estimated a maximum force of ~20 picoNewtons exerted on rat-liver LDs by the optical trap. Because ~80% of fed-LDs escaped from the trap, their binding strength to the fed-mSLB exceeds 20 picoNewtons. In contrast, only ~20% of fasted-LDs trapped above fasted-mSLBs could exceed this force to escape the trap.

Having thus confirmed the physical tethering of LDs to the mSLB, we asked whether LD-mSLB contacts actually result in exchange of lipids and proteins(4, 5, 17). A trace amount of rhodamine-PE (Rh-PE) was incorporated by mixing microsomes with Rh-PE containing liposomes prior to mSLB preparation. RhPE incorporation was similar in fed and fasted mSLBs (Fig S4:A). Next, we added fed-LDs or fasted-LDs to Rh-PE labeled mSLBs. Live imaging showed that fed-LDs acquired rhodamine fluorescence much more rapidly than fasted-LDs (Figs 2:A-D and Supp Movies 3-4). Accordingly, Rh-PE intensity on fed-LDs after 15 minutes of incubation was higher than fasted-LDs (Fig 2E). Protein transfer, a hallmark of inter-organelle communication, has been observed across LD-ER contacts using the model peptide HPos(18). We therefore prepared mSLBs using microsomes from COS7 cells overexpressing mCherry-HPos, incubated them with fed-LDs or fasted-LDs, and imaged LDs after 15 min of incubation (Figs 2F and 2G). Significantly more mcherry-HPos fluorescence could be detected on fed-LDs compared to fasted-LDs (Fig 2H), suggesting efficient transfer of HPos to fed-LDs. MCS formation was also more efficient for fed-LDs, as evidenced from higher *LD_BOUND_* (Fig 2I). Fed-LDs must therefore have factors enabling them to interact with COS-7 mSLBs. We suspected one such factor to be PA because it induces LD-ER interaction in fed state and PA is dramatically enhanced on fed-LDs(8). To test this, we prepared artificial LDs (ALDs)(8) with a phospholipid membrane consisting of PC or PC+PA(=10 mol%), and having similar size (Fig S4:B). Again, more HPos was transferred from COS-7 mSLBs to PA-containing ALDs (Fig 2 J,K,L) and *LD_BOUND_* was higher for PA-containing ALDs (Fig 2M). We further tested if addition of PA to conventional SLBs assists LD-membrane contacts. Both fed-LDs and fasted-LDs showed increased *LD_BOUND_* on PA-containing SLBs as compared to PC-only SLBs (Fig S4:C-G). It is therefore possible that PA, by virtue of its conical shape(19), induces negative curvature to stabilize the neck-like geometry of LD-ER contacts(8).

**Figure 2.**
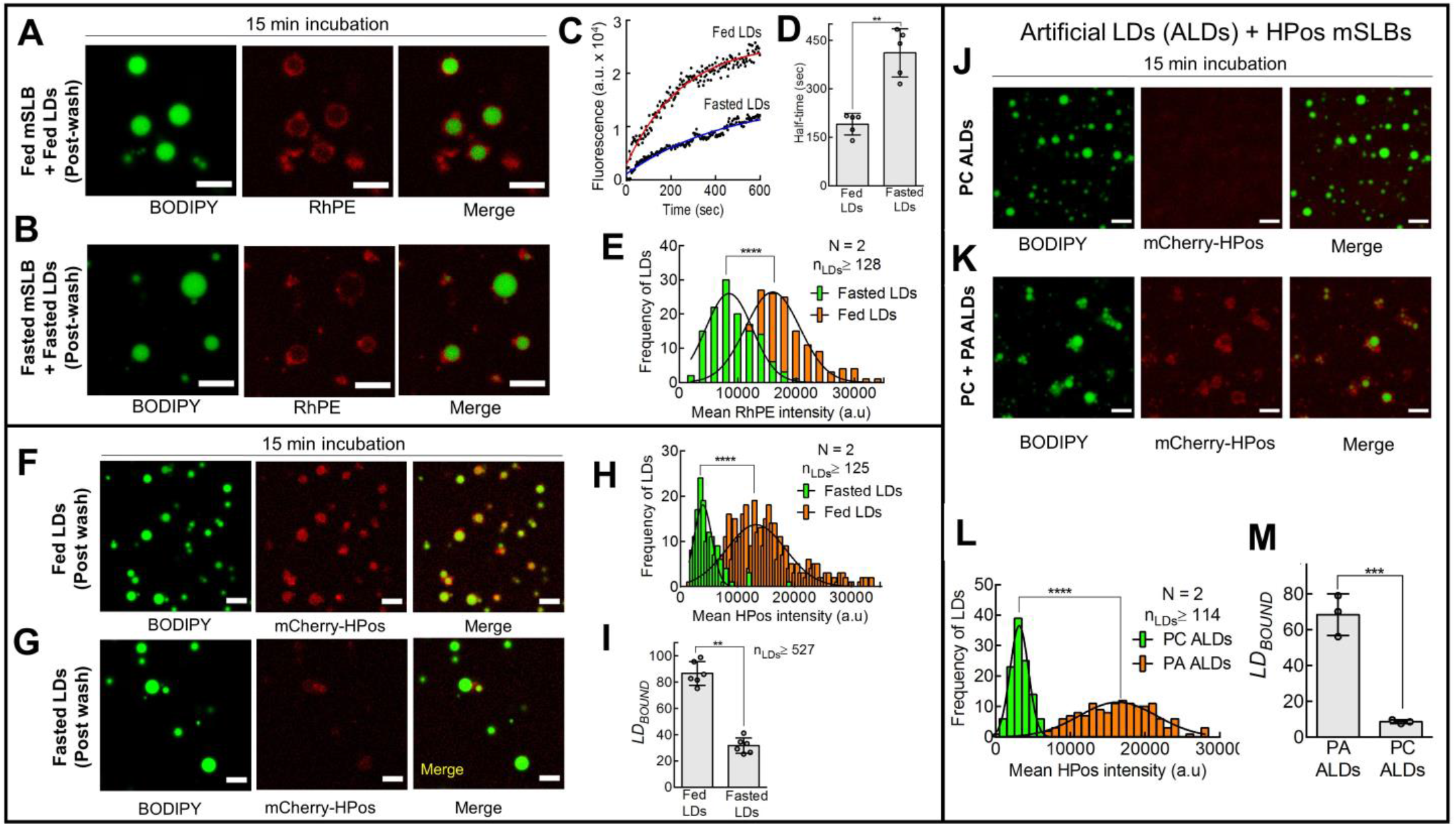
Exchange of lipids and proteins across LD-mSLB contact sites. A. Representative image of fed-LDs incubated for 15 min with fed-mSLB. LDs were labeled with BODIPY (green). The mSLB is doped with rhodamine-PE (RhPE; red). Scale bar = 10 μm. B. Representative image of fasted-LDs incubated with fasted-mSLB for 15 min. Scale bar = 10 μm. C. Change in RhPE fluorescence with time on a single LD of the indicated metabolic state (fed/ fasted) bound to mSLB. D. Half-time for LD fusion with mSLB (as detected by RhPE fluorescence acquisition on LDs). Data represent mean ± S.D. ** *p* = 0.0079, Mann–Whitney test. E. Frequency distribution of RhPE-fluorescence intensity on fed (orange bars, Mean ± S.D. 16005 ± 4657 a.u) and fasted LDs (green bars, Mean ± S.D. 8525± 3996 a.u) after incubation with mSLBs doped with RhPE. Black trace represents a Gaussian fit. N represents the number of independent pair of animals used; n_LDs_ represents the total number of LDs observed. **** *p* < 0.0001, Students t-test. F. Representative image of fed-LDs (green) incubated for 15 min with mSLB (red) made from COS7 cells overexpressing mCherry-HPos. G. Representative images of fasted-LDs (green) incubated for 15 min with mSLB (red) made from mCherry-HPos overexpressing COS7 cells. H. Frequency distribution of mCherry-HPos-fluorescence intensity on fed (orange bars, Mean ± S.D. 13258 ± 5175), and fasted LDs (green bars, Mean ± S.D. 3926 ± 1563) after incubation with mSLBs prepared from COS7 cells overexpressing mCherry-HPos. Black trace represents Gaussian fit. N represents the number of independent pair of animals used; n_LDs_ represent the total number of LDs observed. **** *p* < 0.0001, Students t-test. I. *LD_BOUND_* for fed-LDs and fasted-LDs over mCherry-HPos labeled mSLB prepared from COS7 cells. Data represents mean ± S.D. n_LDs_ represents the total number of LDs observed. ** *p* < 0.01, Mann–Whitney test. J. Representative image of PC-ALDs (green) incubated for 15 min with mCherry-HPos mSLB (red). Scale bar = 10 μm. K. Representative image of PA+PC ALDs (green) incubated for 15 min with mCherry-HPos mSLB (red). Scale bar = 10 μm. L. Frequency distribution of mCherry-HPos-fluorescence intensity on PA (orange bars, Mean ± S.D. 16378 ± 5221 a.u) and PC ALDs (green bars, Mean ± S.D. 3161 ± 1227 a.u) after incubation with mCherry-HPos mSLB. Black line represents a Gaussian fit. *N* represents the number of independent experiments. n_LDs_ represents the total number of LDs observed. **** *p* < 0.0001, Students t-test. M. *LD_BOUND_* for PA+PC ALDs or PC-ALDs incubated with mCherry-HPos labeled mSLB prepared from COS7 cells. Data represent mean ± S.D. *** *p* = 0.0009, Mann–Whitney test.

Our *in-vitro* assay reconstitutes LD-mSLB contacts that exhibit a dependence on metabolic state (fed/fasted) or immune activation(7, 8, 10). Multiple contacts can be observed simultaneously in a single focal plane, making this a high-throughput assay. This planar geometry may also permit incorporation of microfluidics and superresolution TIRF (total internal reflection fluorescence) imaging in future to dissect the molecular architecture of MCS. This assay has limitations because contact between mSLB and coverslip could perturb membrane function, further, asymmetry within the ER bilayer membrane cannot be reproduced on mSLBs. Nevertheless, we could reproduce aspects of LD-ER interactions relevant to fed/fasted transitions(7, 8) and immune activation in cells(10). Further, blocking factors (PA and Rab18) that are important for LD catabolism(8) also inhibited LD-mSLB contacts. The assay could be adapted to interrogate MCS formation between mitochondria, peroxisomes, endosomes, ER and other cellular membranes thus providing much-needed mechanistic insight into neurological, metabolic and pathogenic diseases(3). Such efforts are in progress in our laboratory.

## ACKNOWLEDGEMENTS

We thank S. Ojha and V. Soppina for help and guidance with experiments. N. Vitale for GST-Spo20p construct, A. Pol for HPos construct and R. Parton for Rab18 constructs. S.T. Suryavanshi and Tata Institute of Fundamental Research for maintaining and providing animals. This work was supported by the Department of Atomic Energy – Government of India, Augmented Seed Grant from Indian Institute of Technology Bombay, and Department of Biotechnology – Wellcome Trust India Alliance Fellowships (Grants IA/S/11/2500255 and IA/S/19/2/504634 to R. Mallik).

## AUTHOR CONTRIBUTIONS

SK and RM wrote the manuscript with inputs from other authors. SK, JS and RM designed experiments. SK, JS, HB, ST and MK did the experiments.

## METHODS

### Animal Procedures

Sprague Dawley (SD) rats were bred and maintained by the animal house at Tata Institute of Fundamental Research, Mumbai. All animal protocols were approved by the Institutional Animal Ethics Committee (IAEC) formulated by the Committee for the Purpose of Control and Supervision of Experiments on Animals (CPCSEA), India. Two to three month old male SD rats were used. Rats from the same litter were used for a fed-fasted pair. Rats were maintained in a regular light(12 hour)/dark (12 hour) cycle and fed a standard laboratory chow diet. Fed group rats had *ad libitum* access to food and water. Fasted group rats were fasted for 16 hours, with *ad libitum* access to water. LPS/Saline injections were performed as described(10). Briefly, rats were intraperitoneally injected with 300 μl of saline buffer (control) or 6mg/kg LPS (final dose, dissolved in saline; L2639, Sigma-Aldrich) and fasted overnight (16 hours).

### Isolation of Microsomes

Total rat liver microsomes were isolated using a previously standardized protocol(20). Purity of microsomes was confirmed as described(20) (Fig S1:C). SD rats were used according to following categories:- (i) Fed (ii) Fasted for 16 hours (iii) LPS injected or saline injected [as control], and then fasted for 16 hours. Rats were anesthetized using sodium thiopentane at 40 mg/kg of body weight. Rats were sacrificed and the liver was perfused with 50 ml cold phosphate buffer saline (1XPBS) through the hepatic portal vein. Liver tissue was weighed in 1XPBS and 9gm of liver tissue was placed in 3 volumes of 0.25M sucrose solution having 4mM (DTT), 8ug/ul Pepstatin, 4mM (PMSF) and Roche protease cocktail inhibitor. Mincing and homogenization of tissue was carried out in the cold room. The minced tissue was transferred into 50ml Potter-Elvehjem tissue grinder (kept on ice) and homogenized using ribbed Teflon pestle for up to 20 strokes. The homogenate was filtered through cotton mesh (2 layers) using a glass funnel. The homogenates were centrifuged at 8700*g* at 4°C for 15 minutes to obtain a post nuclear supernatant (PNS). The PNS was spun 43,000*g* (Beckman Coulter ultracentrifuge, Type 70 Ti rotor) at 4°C for 7 minutes to pellet out mitochondria. The microsomes were pelleted at 1,10,000*g* at 4°C for 60 minutes. Microsomes were resuspended in 1XPBS, flash-frozen, and stored at −80°C for further experiments. Microsomes were prepared from cultured cells following similar methods.

### Lipid droplet isolation

Lipid droplets from rat liver were isolated using a previously described protocol^5,6^. Briefly, 2-3 months old male SD rats (in the categories 1,2,3 described above) were anesthetized (sodium thiopentone, 40 mg/kg intraperitoneal injection). The abdominal cavity was cut open to perfuse the liver through the hepatic portal vein with cold PBS. The perfused liver was dissected, washed and weighed. The liver was minced and resuspended in 1.5 times wt/vol of 0.9M sucrose containing MEPS buffer, and homogenized using Dounce homogenizer at 4°C. MEPS buffer is composed of 35 mM PIPES, 5 mM EGTA, and 5 mM MgSO_4_ pH 7.1, supplemented with protease inhibitor cocktail (Roche), 4mM PMSF (Sigma), 8 μg/ml pepstatin A (Sigma) and 4 mM DTT (Sigma). We centrifuged the homogenate at 1,800 g for 10 minutes at 4°C to obtain PNS. The PNS thus obtained was mixed with 1.5 times vol/vol of 2.5 M sucrose containing MEPS buffer (without PMSF) and was loaded as the bottom layer of sucrose density gradient. This layer was overlaid with 5 ml (each) of 1.4M, 1.2M, 0.5M, and 0M sucrose in MEPS buffer. This gradient was centrifuged at 120,000 g at 4°C for 1 hour to obtain lipid droplets (top-most whitish layer). LDs were collected using an 18G needle, flash-frozen, and stored in liquid nitrogen.

### Preparation of Liposomes and Small unilamellar vesicles (SUVs)

A mixture of DOPC, EggPA and RhPE (Avanti Polar Lipids) in the ratio of 89.5:10:0.5 mol% for PA SLB and 99.5:0:0.5 mol% for PC SLB lipids was aliquoted in a glass test tube (final concentration of 3mM) and mixed gently. This chloroform dissolved lipid mix was dried rapidly under a nitrogen stream, and vacuum desiccated for 60 minutes. The dried lipid film was hydrated in 1XPBS at 50°C for 30 minutes. After incubation, the hydrated liposomes were vigorously vortexed for 5 minutes. The hydrated liposomes were sonicated using a probe sonicator for 5 minutes to make SUVs (Branson SFX250 Sonifier^®^; 15% Amplitude, 3sec ON and 2sec OFF cycle). The sample was centrifuged at 20,000g for 10min, 4 degC to remove LUVs (large unilamellar vesicles) and titanium particles (eroded from the probe).

### Conventional SLB and Microsomal SLB (mSLB) preparation

The procedure is adapted from earlier protocols(11). For conventional SLBs, a glass coverslip was cleaned by piranha [6:4 vol/vol concentrated H_2_SO_4_: H_2_O_2_] treatment or by oxygen plasma (Harrick plasma PDC-32G). The coverslip was then assembled in a flow cell using double-sided tape. SUVs were introduced into the flow cell and allowed to fuse with the hydrophilic surface of the coverslip for 45 minutes inside a humid chamber. After incubation excess unbound liposomes were washed off using 1XPBS (200μl x 3 times).

For mSLBs, the flowcell was prepared as above. Total rat liver microsomes (protein concentration 5 mg/ml) were sonicated for 5 minutes using a probe sonicator (Branson SFX250 Sonifier^®^; 15% Amplitude, 3sec ON and 2sec OFF cycle). The sample was centrifuged at 20,000g for 10 minutes to remove aggregated proteins. These sonicated microsomes (protein concentration 1.5 mg/ml) were labeled with BODIPY-C12 (Invitrogen™, product number: D3822, 8μM final probe concentration). Alternatively, microsomes were mixed with sonicated SUVs made from DOPC:RhPE liposomes for labelling (DOPC:RhPE 99.5:0.5 mol%, 0.2mM final concentration). Microsomes were then introduced into the flowcell and allowed to deposit on the cleaned coverslip for 45 minutes in a humid chamber. After incubation, excess unbound microsomes were washed off using 1XPBS (200μl X 3 washes) and the sample was imaged. LDs isolated from rat liver were then introduced into the chamber and allowed to form contacts for 15 minutes. The flowcell was kept in a humid chamber during this period. The sample was imaged to validate normalization of LDs (before washing) and then washed off with an equal volume of buffer. The contacts between LDs and mSLB were imaged in a confocal microscope.

### Preparation of Artificial Lipid Droplets (ALDs)

ALDs were prepared by adopting a previously described using a freeze-thaw technique(8). Briefly, 70 μl glyceryl trioleate (TG) and 0.5 μmol egg PC were mixed in acid-washed clean glass tubes to prepare 1 ml of ALDs. We made PA-ALDs by supplementing the above reaction mix with 25nmol of Egg-PA. This mix was also supplemented with trace amounts of rhodamine-PE (1.6 nmol) to aid imaging and detect lipid exchange. This reaction mixture was dried under nitrogen gas stream for 30 minutes followed by vacuum desiccation for 1 hour to remove trace amounts of chloroform. After desiccation, 930μl HKM buffer (50 mM HEPES-KOH, 120 mM potassium acetate, and 1 mM MgCl_2_, at pH 7.4) was added to the reaction mix. This reaction mixture was vigorously vortexed for 10 minutes to aid emulsification. This whitish emulsion was subjected to five freeze-thaw cycles of flash freezing in liquid N2 and thawing at 55°C. Each cycle was accompanied by vigorous vortexing for 10 minutes. ALDs were stored in liquid nitrogen and thawed prior to experiments.

### Antibody inhibition experiment

mSLBs were prepared as described above and incubated with relevant antibody for 30 minutes in a humid chamber. After incubation, excess antibody was washed off from mSLBs, LDs were introduced and incubated for 15 minutes. After incubation, the unbound/weakly bound LDs were washed off and stable contacts LD-mSLB contacts were imaged.

### Fluorescent Recovery after Photobleaching (FRAP)

FRAP analysis was performed on mSLB/conventional SLB labeled with BODIPY-C12. The total scan area (2025μm^2^) and bleached region (6μm radius) was restricted to obtain high temporal resolution. Images were acquired at 1x optical magnification. Argon laser (488nm) at 8% power and 150 iterations at 100% transmission were used for photobleaching. Standardization of imaging parameters was performed to minimize photobleaching during acquisition. The mean post-bleached intensity was normalized to the mean pre-bleach intensity. The recovery kinetics (characteristic diffusion time) was analyzed by fitting data to the recovery equation Y=(F_0_+F_i_*(X/t_half_))/(1+(X/t_half_)) as described(21).

### Plasmids

GST-Spo20p (PABD) was a gift from Nicolas Vitale (Universit’e de Strasbourg, France). HPos overexpression construct was a gift from Albert Pol (Universitat de Barcelona, Spain). GFP-Rab18 (Q67L) and GFP-Rab18 (S22N) was a kind gift from R.G Parton (University of Queensland, Australia).

### Antibodies

ADRP antibody (651102) was purchased from PROGEN Biotechnik. Hsp60 antibody (ab46798) was purchased from Abcam. PDI antibody (612117) was purchased from BD Bioscience. Anti-FLAG antibody (F1804) was purchased form Sigma. Rab18 antibody (sc393168) and Calnexin (sc 11397) antibody were purchased from Santa Cruz Biotechnology. S6 ribosomal protein antibody (2317) was brought from Cell Signaling Technologies. Alexa-594 labeled anti-mouse and anti-rabbit secondary antibodies were purchased from Invitrogen.

### Cell Culture, transfection and Microsome preparation

COS-7 (ATCC, CRL-1651)/HEK293T cells were maintained in DMEM containing 10% FBS. Six 100-mm plates of COS-7 or HEK293T cells were transfected with mCherry-HPos or Rab18 plasmids. Microsome isolation from cells was performed as described(7, 8). Briefly, cells were resuspended in 3 ml hypotonic buffer (10 mM HEPES pH 7.4, 1mM EDTA supplemented with protease inhibitor cocktail) and incubated for 30 minutes (4°C). After incubation, the cells were centrifuged at *1200g* for 10 minutes, and the cell pellet was resuspended in isotonic buffer (250 mM sucrose, 10 mM HEPES pH 7.4, 1 mM EDTA, protease inhibitor cocktail). Cell lysis was performed in a cell cracker (Isobiotec;14-micron clearance; 10 strokes). The lysate was centrifuged at 1500*g* for 10 minutes to pellet cell debris. The resulting supernatant was centrifuged again at 12,000*g* for 15 minutes to pellet mitochondria. The resulting supernatant was fractionated into LDs, soluble, and membrane fractions (enriched in microsomes) by using a two-layer sucrose gradient. The bottom layer consisted of 1.5 ml of 250 mM sucrose, 10mM Hepes pH 7.4, 1 mM EDTA, and a complete protease inhibitor tablet (Roche). The upper layer consisted of 3 ml of 50 mM sucrose, 10 mM Hepes pH 7.4, 1 mM EDTA, and a complete protease inhibitor tablet. Centrifugation was done at 100,000*g* for 6hrs. Protein concentration was determined using the BCA method (Sigma).

### Optical trapping

The instrument has been described(13), briefly LD were observed at room temperature in a custom developed DIC microscope (Nikon TE2000-U) using a 100x, 1.4 NA oil objective. Image frames were acquired at video rate (30 frames/sec) with a Cohu 4910 camera. Individual LDs were captured in the optical trap and lowered to be brought in contact with the mSLB surface. A piezo-stage was then used to move the mSLB below the trapped LD at a constant velocity of 1 μm/sec. The position of LDs was tracked frame-by-frame using custom written software(22).

### GST-tagged Spo20p purification

The GST fusion proteins were expressed in bacteria *(E. coli,* BL21) and purified by glutathione-Sepharose (Cat No. 27-4574-01, Qiagen) as per the manufacturer’s instructions. The purified proteins were dialyzed against 1XPBS. The samples were checked for quality by running on SDS-PAGE followed by Coomassie Brilliant Blue staining.

### Thin Layer Chromatography

Lipids from LDs were extracted using chloroform extraction^5,6^. Briefly, 0.8 ml aqueous sample containing LDs was mixed with 2ml of methanol and 1ml chloroform followed by vortexing. 1ml each of chloroform and water was added, which resulted in phase separation. The lower organic phase was transferred to a new glass tube, dried under a N_2_ stream, and resuspended in 20μl of chloroform. The silica TLC plates (Merck) were precleaned using chloroform followed by air-drying. The sample was then spotted onto these plates using a glass capillary. The solvent system used was according to Wilfling and Thiam et al.(23) with minor modifications. The first solvent system, containing a mixture of n-hexane/diethyl ether/acetic acid (70:30:1), was run halfway and air-dried. The plate was then run in a solvent mixture of n-hexene/diethyl ether (59:1). The plate was dried and visualized by spraying with 10% CuSO4 in 8% H_3_PO_4_ followed by baking in the oven above 150°C for 15-20 minutes.

### Antibody staining of mSLBs

mSLBs were incubated with an equal concentration of respective primary antibodies and incubated overnight at 4°C in a humid chamber. After incubation primary antibody was washed off using 1XPBS and mSLB was incubated with fluorescently labeled secondary antibody (1:200 dilution) for 1 hour. The excess unbound secondary antibody was then washed off and the sample was imaged.

### Microscopy

mSLBs and LDs were imaged using a Zeiss LSM 880 upright laser scanning microscope equipped with a 63X, 1.4 NA oil-immersion objective. The sequential excitation of fluorophores was achieved by using 488 nm and 561 nm lasers. Spectral bandpass emission filters were used for the acquisition of confocal images with high-sensitivity detectors. Imaging parameters were kept constant across the compared experiments.

### Immunoblotting

Equal concentration of protein samples were resolved on a 10% SDS-PAGE gel transferred to PVDF membrane for immunoblotting. The blot was blocked for 1 hour at room temperature with 5% non-fat dry milk in Trisbuffered saline with 0.1% Tween-20 (blocking buffer). The membrane was incubated with primary antibody diluted in the blocking buffer for 1 hour at room temperature or overnight at 4°C. The membrane was washed with 0.1% TBST three times for 15 minutes each. The membrane was incubated with HRP-conjugated secondary antibody diluted in the blocking buffer for 1 hour at room temperature. The membrane was washed with 0.1% TBST three times for 15 minutes each and developed using ECL kit (Millipore). The blots were imaged on an Amersham Imager 600 and band intensity was quantified using Image J.

### Image and statistical analysis

Image analysis was carried out using ImageJ-Fiji. Statistical analysis was performed using GraphPad Prism (version 5.0a). n_LDS_ refers to the number of LDs and tubes analyzed in a single experiment. *N* refers to the number of independent experiments.

---

**Supplementary Figure 1.**
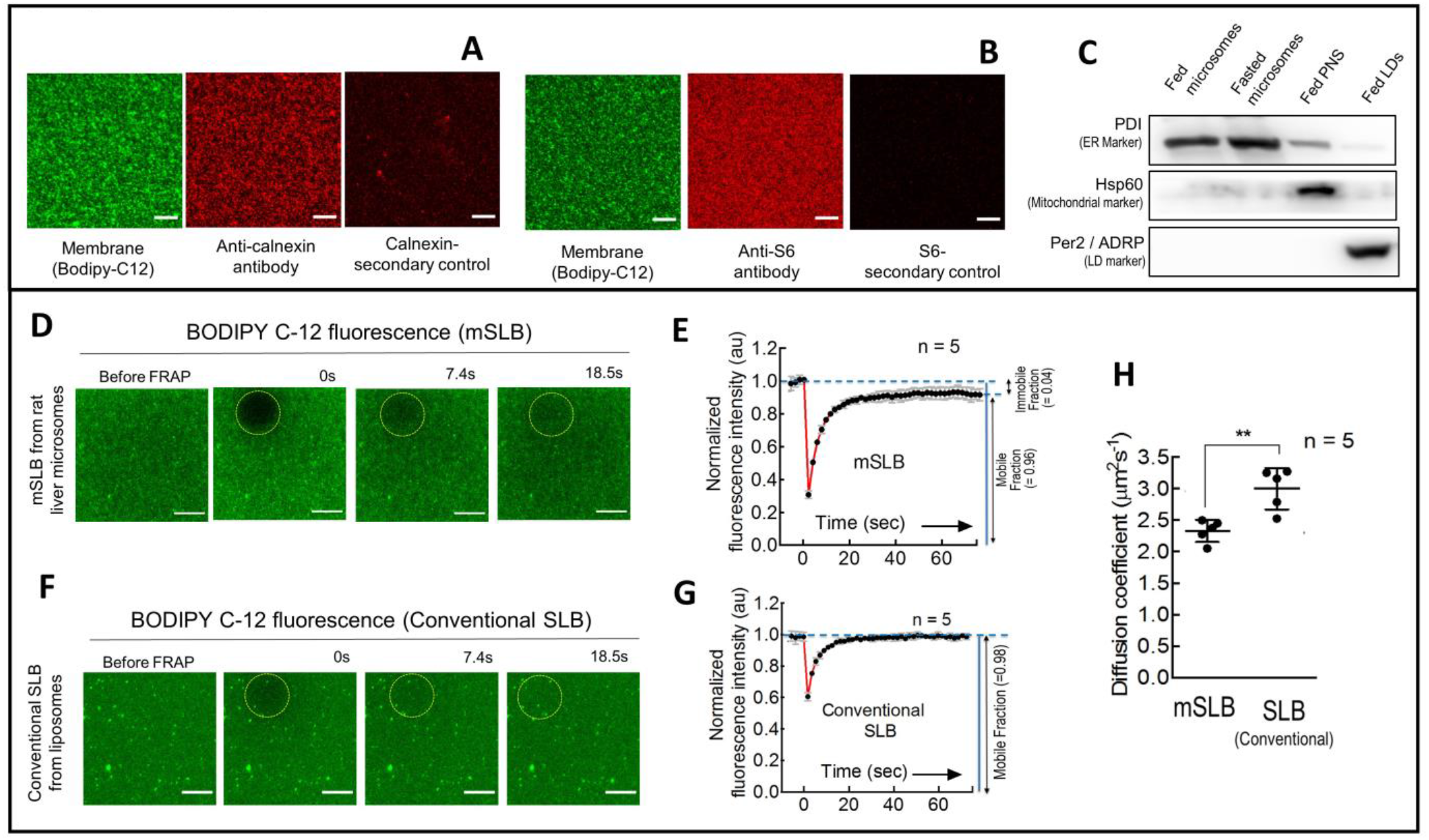
Validation of BODIPY-C12 as a membrane marker and related experiments. A. Representative image of mSLB labeled with BODIPY-C12 (green). The ER marker calnexin was detected using a calnexin-specific primary antibody and a secondary antibody (Red). Images with and without primary antibody are shown. Scale bar = 10 μm. B. Representative image of mSLB labeled with BODIPY-C12 (green). The ER marker S-6 was detected using an S-6 specific primary antibody and a secondary antibody (Red). Images with and without primary antibody are shown. Scale bar = 10 μm. C. Immunoblot showing purity of microsomes and LDs. Fed or fasted microsomes, fed post nuclear supernatant and fed lipid droplets were loaded with equal protein content in all samples. Western blot was done using antibodies against PDI (ER marker), HSP 60 (mitochondria marker) and ADRP (LD marker). D. Fluorescence Recovery after Photobleaching (FRAP) of BODIPY-C12 labeled mSLB prepared using microsomes from liver of fed rat. Dotted circle marks the bleached region. Scale bar = 10 μm. E. Fluorescence recovery data of mSLBs fitted to a recovery equation [red trace; Y=(F_0_+F_i_*(X/T_half_))/(1+(X/T_half_))]. Black dots represent mean and error bars represent S.D. Data is average of 5 different experiments. High mobility of lipids within the mSLB is inferred from the high mobile fraction observed (see definition in figure). F. FRAP of BODIPY-C12 labeled conventional SLB. The SLB was prepared using DOPC:EggPA liposomes {89:10 mol%}. The dotted circle marks the bleached region. Scale bar = 10 μm. G. Fluorescence recovery data of conventional SLBs fitted to recovery equation [red trace]. Black dots represent mean and error bars represent S.D. Data is average of 5 different experiments. High mobility of lipids within the mSLB is inferred from the high mobile fraction observed. H. Calculated diffusion coefficients for BODIPY-C12 labelled conventional SLB and mSLB. Five different recovery curves for each case were fitted to the recovery equation to calculate half-time and diffusion coefficient of bleached lipids.

**Supplementary Figure 2.**
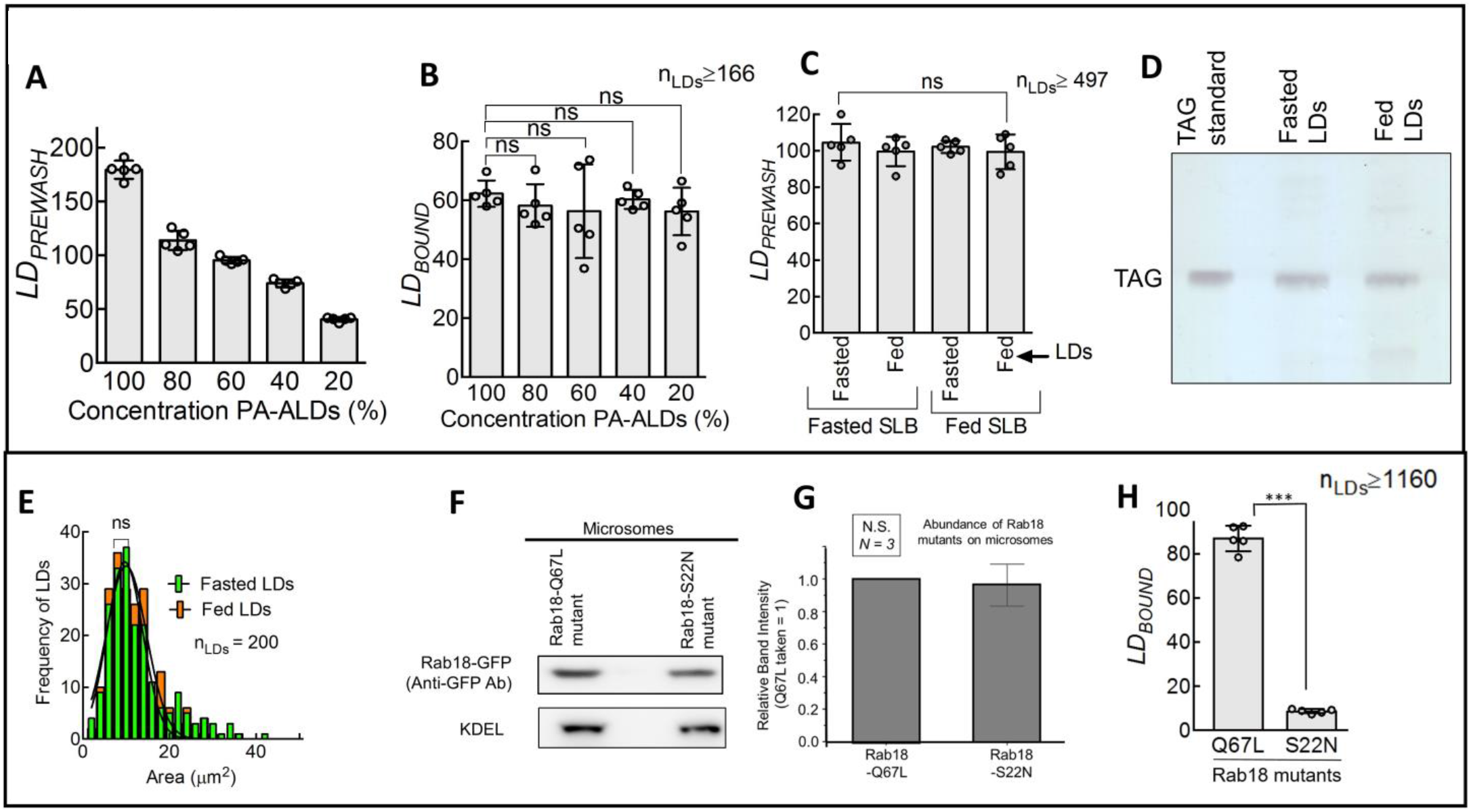
Quantitation of LDs before washing and related experiments. A. Quantitation of prewash PA-ALDs under different dilutions. Each point on the graph represents a field of view. Data represent mean±SD. B. Quantitation of *LD_BOUND_* under different PA-ALDs concentration. Each point on the graph represents a field of view. Data represent mean ± SD. n_LDs_ represents the total number of LDs observed. ns = not significant, Mann–Whitney test. C. *LD_PREWASH_* (see main text) is same for LDs across different metabolic states. Each data point on the graph represents a field with ~80-100 LDs. Data represent mean ± S.D. ns = not significant, Mann–Whitney test. D. Thin Layer Chromatography (TLC) to detect triglyceride (TAG) in fed-LD and fasted-LD samples after normalization using OD_600_. E. Frequency distribution of the area of individual fed-LDs (orange bars, Mean ± S.D. 10.0 ± 4.6 μm^2^) and fasted-LDs (green bars, Mean ± S.D. 9.6 ± 4.1 μm^2^). Black trace is a Gaussian fit. n_LDs_ represents the total number of LDs observed. ns = not significant, Students t-test. F. Immunoblot for detecting Rab-18 mutants on microsomes. Microsomes isolated from Rab18 Q67L and S22N mutant HEK293T cells (having equal protein content) were loaded and probed using anti-GFP antibody. KDEL was used as loading control. G. Quantitation of western performed in F. Anti-GFP intensity is normalized to KDEL signal. *N* represents the number of independent experiments. H. *LD_BOUND_* calculated for fed-LDs to mSLBs overexpressing different Rab18 mutants. Each point on the graph represents a field of view with ~100 LDs. Data represents mean ± S.D. n_LDs_ represents the total number of LDs observed. *** *p* < 0.001, Mann–Whitney test.

**Supplementary Figure 3.**
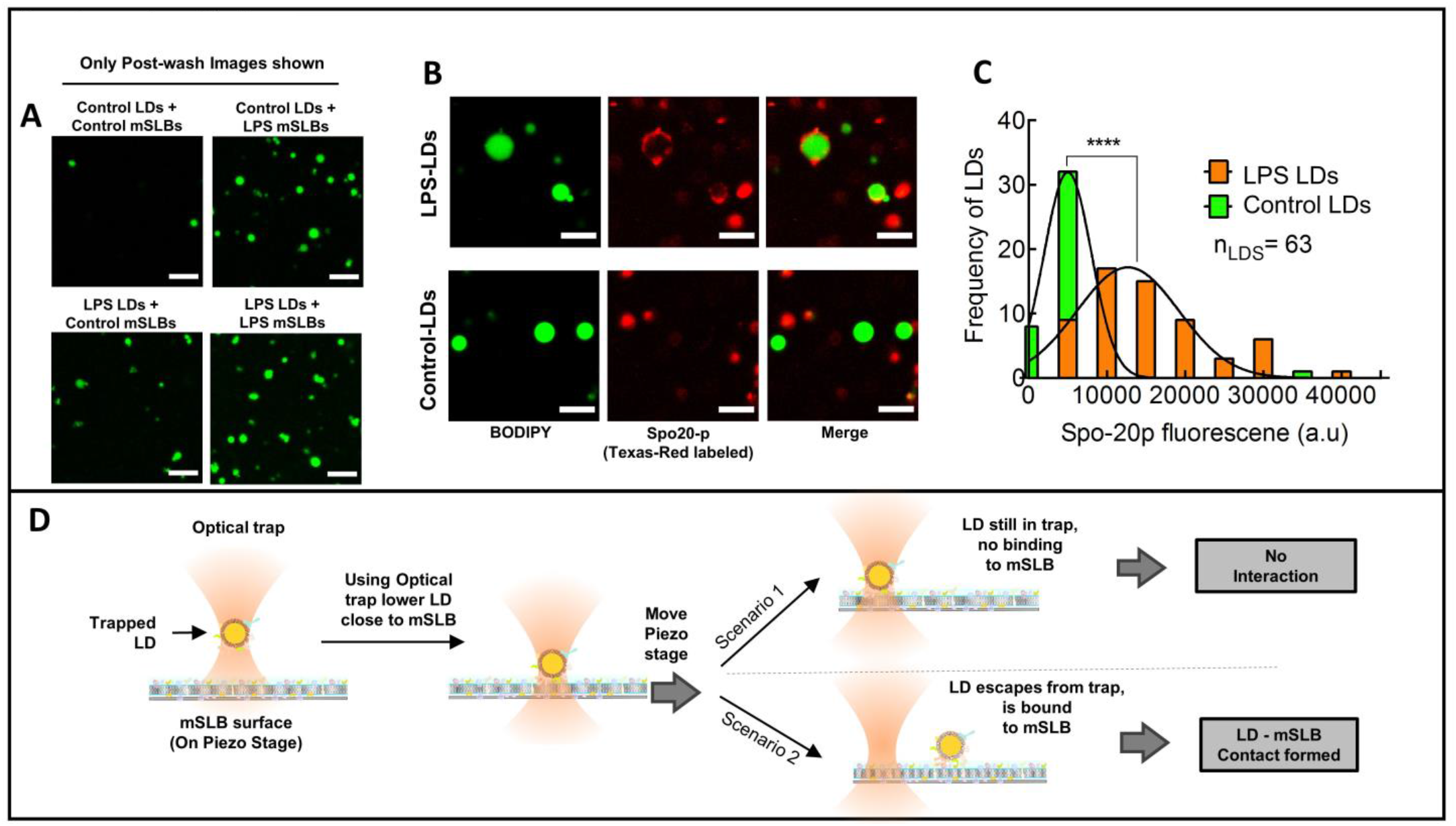
LPS activation promotes stable LD-mSLB contacts. A. Representative fluorescence micrographs of LDs isolated from control (saline injected fasted) and LPS (LPS injected fasted) rat liver. LDs are labeled with BODIPY (green). The LDs were incubated with mSLBs prepared from control (saline injected fasted) or LPS (LPS injected fasted) rats and imaged (Pre-wash; images not shown). The flowcell was washed and LDs imaged again (Post-wash; shown in figure). *LD_BOUND_* was calculated as described. Scale bar = 10 μm. B. Representative fluorescence micrographs of Texas-Red maleimide^®^ labeled-Spo20p binding to LDs isolated from control and LPS treated rat liver. Images are taken 15-min post incubation. The LDs were incubated with labeled-Spo20p prepared from fasted control or LPS injected rats. LDs labeled with BODIPY are in green. Spo20p is fluorescently tagged with Texas-Red^®^ C-2 maleimide is in red. Scale bar = 5 μm. C. Frequency distribution of Spo20p fluorescence intensity on LPS treated (orange bars, Mean ± S.D. 12700 ± 6521), and Control LDs (green bars, Mean ± S.D. 5014 ± 3022). Black trace represents Gaussian fit. n_LDs_ represent the total number of LDs observed. **** *p* < 0.0001, Students t-test. D. Schematic representing tethering assay using mSLB. A single LD is optically trapped and brought in contact with mSLB surface. The entire flow cell is displaced (keeping the trap position invariant) using a piezo-stage with a constant velocity of 1 micron/sec. If the LD does not bind strongly to the mSLB surface, then it remains in the trap. If LD binds strongly to the mSLB surface (by forming MCS), then it escapes from the trap.

**Supplementary Figure 4.**
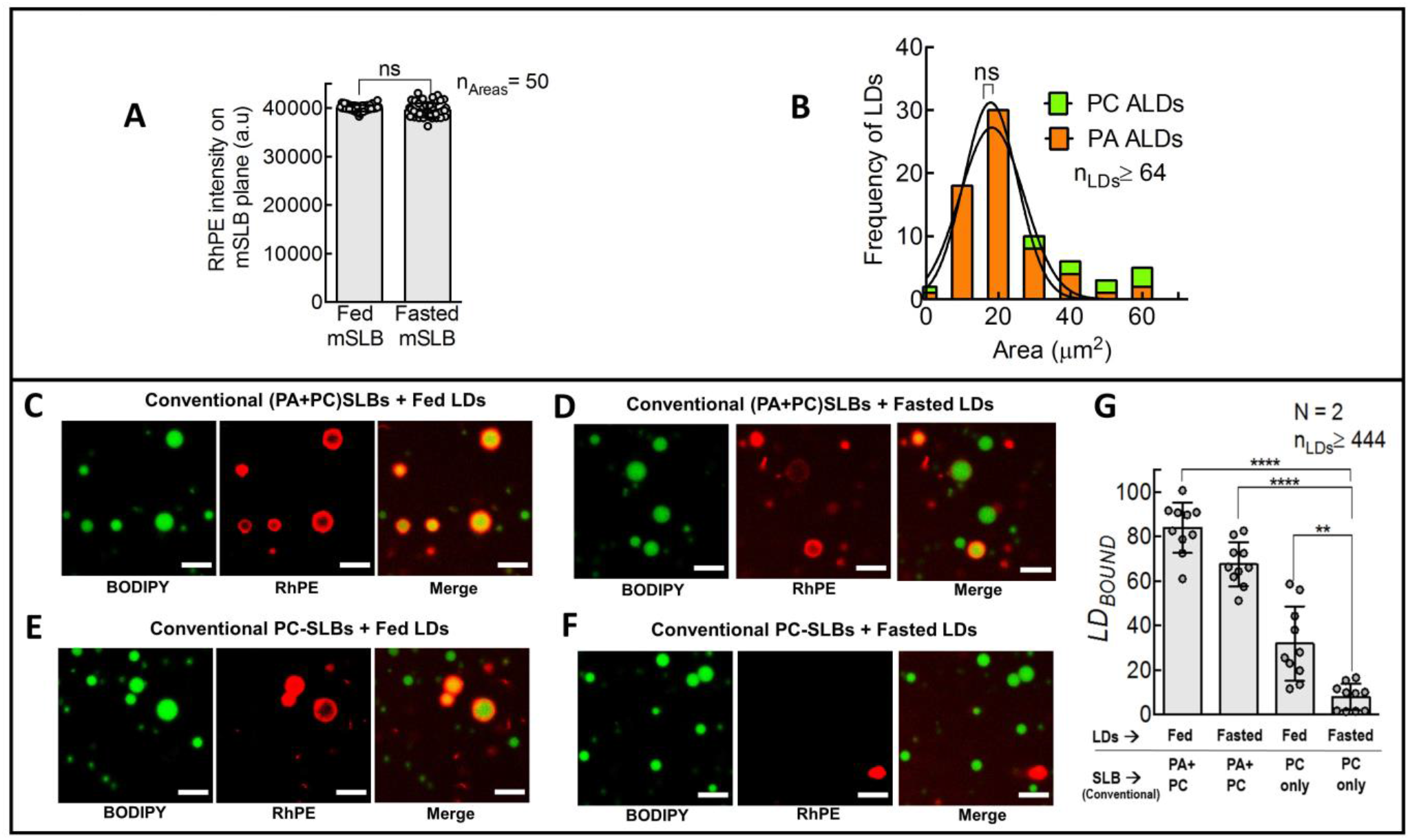
Role of Phosphatidic acid in LD-mSLB contacts and related experiments. A. Plot comparing RhPE fluorescence on the fed and fasted mSLB surface. RhPE is introduced into mSLB by incubating microsomes with liposomes containing DOPC:RhPE (99.5:0.5 mol%). The area used for calculating fluorescence intensity was same. Data represent mean ± S.D. Fifty fields-of-view were analyzed for each condition. ns = not significant, Mann-Whitney test. B. Frequency distribution of area of ALDs prepared using PC+PA (orange bars, Mean ± S.D. 17.8 ± 7.4 μm^2^) and PC (green bars, Mean ± S.D. 18.2 ± 8.8 μm^2^). Black trace is a Gaussian fit. n_LDs_ represents the total number of LDs observed. ns = not significant, Students t-test C. Fluorescence micrograph of fed LDs (BODIPY labeled, green) incubated with conventional SLB containing PC and PA (red; prepared by fusion of liposomes having DOPC:EggPA:RhPE 89.5:10:0.5 mol%). D. Fluorescence micrograph of fasted LDs (BODIPY labeled, green) incubated with conventional SLB containing PC and PA (red; prepared by fusion of liposomes having DOPC:EggPA:RhPE 89.5:10:0.5 mol%). E. Fluorescence micrograph of fed LDs (BODIPY labeled, green) incubated on conventional SLB containing PC (red; prepared by fusion of liposomes having DOPC:RhPE 99.5:0.5 mol%). F. Fluorescence micrograph of fasted LDs (BODIPY labeled, green) incubated on conventional SLB containing PC (red; prepared by fusion of liposomes having DOPC:RhPE 99.5:0.5 mol%). G. Quantitation of *LD_BOUND_* for LDs (isolated from the liver of fed rat) on conventional SLBs. Each data point on the graph represents a field of view with ~100 LDs across multiple experiments. Data represents mean ± S.D. **** *p* < 0.0001, ** *p* = 0.003, Mann–Whitney test.

## MOVIE CAPTIONS

**Supplementary Movie 1. Tethering of fed LD on fed mSLB using optical trap**

A fed-LD is optically trapped and brought in contact with fed mSLB. Piezo-stage is then moved with constant velocity of 1 μm/sec to gauge interaction. Scale bar = 5 μm.

**Supplementary Movie 2. Tethering of fasted LD on fasted mSLB using optical trap**

Fasted LD is optically trapped and brought in contact with fasted mSLB. Piezo-stage is then moved with constant velocity of 1 μm/sec to gauge interaction. Scale bar = 5 μm.

**Supplementary Movie 3. Real-time imaging of fed LDs making contacts with fed mSLBs**

Merged movie of fed LDs (Green, labeled with BODIPY) with fed mSLB (Red, doped with RhPE labeled liposomes). Scale bar = 10 μm.

**Supplementary Movie 4. Real-time imaging of fasted LDs making contacts with fasted mSLBs**

Merged movie of fasted LDs (Green, labeled with BODIPY) with fasted mSLB (Red, doped with RhPE labeled liposomes). Scale bar = 10 μm.

